# *Aspergillus nidulans* cell wall integrity kinase, MpkA, impacts cellular phenotypes that alter mycelial-material mechanical properties

**DOI:** 10.1101/2024.09.28.615565

**Authors:** Kelsey Gray, Harley Edwards, Alexander G. Doan, Walker Huso, JungHun Lee, Wanwei Pan, Nelanne Bolima, Meredith E. Morse, Sarah Yoda, Isha Gautam, Steven D. Harris, Marc Zupan, Tuo Wang, Tagide deCarvalho, Mark R. Marten

## Abstract

Mycelial materials are an emerging, natural material made from filamentous fungi that have the potential to replace unsustainable materials used in numerous commercial applications (e.g., packaging, textiles, construction). Efforts to change the mechanical properties of mycelial-materials have typically involved altering growth medium, processing approaches, or fungal species. Although these efforts have shown varying levels of success, all approaches have shown there is a strong correlation between phenotype (of both fungal mycelia and mycelial material’s assembly) and resultant mechanical properties. We hypothesize that genetic means can be used to generate specific fungal phenotypes, leading to mycelial materials with specific mechanical properties. To begin to test this hypothesis, we used a mutant of the model filamentous fungus, *Aspergillus nidulans*, with a deletion of the last kinase in the cell wall integrity (CWI) signaling pathway, mpkA. We generated one set of mycelial materials from the Δ*mpkA* deletion mutant (A1404), and another from its isogenic parent (A1405; control). When subjected to tensile testing, and compared to material generated from the control, Δ*mpkA* material has similar elastic modulus, but significantly increased ultimate tensile strength, and strain at failure. When subjected to a fragmentation assay (i.e., resistance to shear-stress), the Δ*mpkA* material also had higher relative mechanical strength. To determine possible causes for this behavior, we carried out a comprehensive set of phenotype assessments focused on: three-dimensional structure, hyphal morphology, hyphal growth behaviors, and conidial development. We find, compared to the control, material generated from the Δ*mpkA* mutant manifests significantly less development, a modified cell wall composition, larger diameter hyphae, more total biomass, higher water capacity and more densely packed material, which all appear to impact the altered mechanical properties.

## 1. Introduction

An accumulation of societal pressures, such as increased populations, heightened consumption of products, escalated use of toxic additives, and the depletion of non-renewable resources, has encouraged the exploration of sustainable materials to maintain (or even advance) our way of living [1-3]. Fully biobased materials have the potential to alleviate these pressures due to their 100% renewability and their ability to decompose naturally [1, 4, 5]. Furthermore, nature has already demonstrated its aptitude for producing materials that possess some of the most elite mechanical properties known to man (e.g., silk’s strength rivals steel and Kevlar) [6]. Unlocking the full potential of various biobased materials will require a stronger understanding of how the composition, assembly, and growth behaviors of these materials impact resultant mechanical properties. Knowledge of these factors will allow manipulation of these traits to achieve specific and desirable mechanical properties [5].

Materials made out of filamentous fungi, or mycelial materials, represent one of the most promising fully-biobased materials available [7]. Mycelial materials have been championed due to their high mechanical strength, ability to be easily composted, inexpensive production costs, and tolerance to various environments [8, 9]. Mycelial materials are typically produced in one of two forms [10]. Composites consist of a fungal-substrate mixture (e.g., fungi and sawdust), while pure materials are made exclusively from fungi, typically grown in liquid medium. In both cases, the specific fungus used (i.e., species and strain) contributes significantly to the resultant material’s mechanical properties [9, 11]. Composite mycelial materials offer both advantages and disadvantages related to the mechanical properties of the substrates used [12]. In contrast, pure mycelial materials offer the advantage of increased freedom in how they can be produced and utilized [13]. This freedom stems from the pure mycelial material’s independence of other substances that can influence cost, application limitations, mechanical properties, and other aspects that impact pure mycelial materials usefulness [13].

As a result of this flexibility, pure mycelial materials are being used, or considered, for a broad range of applications. For example, they are already being used as replacements for foam and leather and may eventually replace polystyrene packaging in addition to being used as support material for electronics [13]. There is also potential for pure mycelial materials to take on identities as building materials (e.g., sound and heat insulation), nanopapers [12, 14], and engineered living materials [15]. However, for a number of applications, the mechanical demands exceed the capacity of pure mycelial materials, and thus there is a need for improvement in mycelial material mechanical properties [13, 16]. Past reports of improvements in the mechanical properties of pure mycelial materials (e.g., *Ganoderma lucidum* or *Pleurotus ostreatus*) have typically involved changing growth medium [17]. Others have treated mycelial materials with plasticizers, increasing the material’s elasticity, but reducing its ultimate tensile strength [18, 19]. An exciting opportunity to modify the mechanical properties of mycelial materials lies in genetic manipulation of the fungi involved. However, few reports exist using this approach [20, 21].

Two recent studies have explored the idea of evaluating fungal phenotypes as drivers for differences in mechanical properties of resultant mycelial materials [17, 21]. In these studies, scanning electron microscopy (SEM) was used to characterize mycelial-material packing density and surface morphology [17, 21]. Atomic force microscopy (AFM) was used to characterize the dimensions of hyphae in the z-axis, in addition to mechanically testing mycelial materials on a microscale [17]. FTIR was used for collecting data regarding the chemical composition of each fungus tested [17, 21]. The information gathered through these assessments implied a strong connection between the measured phenotypes and resultant material mechanical properties.

While some phenotype-material connections have been made, it has been suggested that more robust characterization of fungal phenotypes (e.g., improved understanding of morphologies), and how they impact material mechanical properties, is required to fully realize the potential of mycelial materials [13, 16]. To this end, our overarching hypothesis is that genetic means can be used to rationally tune fungal phenotype, resulting in mycelial materials with specific mechanical properties. To begin to test this hypothesis, we used a mutant of the model fungus *Aspergillus nidulans* with a deletion of the last kinase in the cell wall integrity (CWI) signaling pathway, *mpkA*. The CWI pathway is responsible for cell wall integrity and repair [22] and is highly conserved in the fungal kingdom [23]. Deletion of *mpkA* (Δ*mpkA*) renders the CWI pathway inoperative [24], and leads to significant morphological and growth defects [25]. Accordingly, we initially hypothesized that an inoperative CWI pathway would lead to weaker-walled hyphae and result in mycelial material with reduced ultimate tensile strength. To test this hypothesis, we grew mycelial materials from both the Δ*mpkA* mutant and its isogenic parent (control). We characterized the mechanical properties of the resultant mycelial material and find the Δ*mpkA* mutant produces material with increased ultimate tensile strength, strain at failure, and resistance to shear-stress. To understand why this occurs, we carried out an extensive set of assessments to better understand how this deletion impacts the phenotype of the mutant, and its corresponding material.

## 2. MATERIALS AND METHODS

### 2.1 Fungal Strains and Growth Media

All experiments were carried out using a mutant containing an *mpkA* deletion (ΔmpkA; FGSC 1404; *pyrG*^*AF*^ used to replace *mpkA*) and an isogenic control (FGSC 1405; *pyrG* transformed back into SO451) [26] obtained from the Fungal Genetic Stock Center (FGSC, Manhattan, KS) [27]. Both genotypes were generated from the SO451 background (*pyrG89; wA3; argB2;* Δ*nkuAku70::argB pyroA4; sE15 nirA14 chaA1 fwA1*) [28]. Both the ΔmpkA mutant and the control were first cultivated on modified MAG-V agar plates containing 2 g/L BD Bacto Peptone, 1 mL/L vitamin solution, 1 mL/L Hutner’s trace elements solution, 20 g/L granulated agar, 1.12 g/L uracil, 1.22 g/L uridine, 35.065 g/L NaCl, 20 g/L glucose, and 20 g/L malt extract. The trace element solution was made of: 22 g/L ZnSO_4_·7H_2_O, 11 g/L H_3_BO_3_, 5 g/L MnCl_2_·4H_2_O, 5 g/L FeSO_4_·7H_2_O, 1.7 g/L CoCl_2_·6H_2_O, 1.6 g/L CuSO_4_·5H_2_O, 1.5 g/L Na_2_MoO_4_.2H_2_O and 50 g/L EDTA (Na_4_). The vitamin solution contained: 100 mg/L biotin, 100 mg/L pyridoxine, 100 mg/L thiamine, 100 mg/L riboflavin, 100 mg/L p-aminobenzoic acid, and 100 mg/L nicotinic acid [29, 30]. Modified YGV was used to grow mycelial materials. 1 L of modified YGV media contained 1 g/L yeast extract, 2 g/L BD Bacto peptone, 1 g/L Bacto casamino acids, 1 mL/L vitamin solution, 1 mL/L Hutner’s trace elements solution, 44.21 g/L KCl, 1.12 g/L uracil, 1.22 g/L uridine, 20 g/L glucose, 20 g/L malt extract, 10 g/L proline, 50 mL/L nitrate salts, and 5 mL/L MgSO_4_ solution. The nitrate salts are made from 142.7 g/L KNO_3_, 10.4 g/L KCl, 16.3 g/L KH_2_PO_4_, and 20.9 g/L K_2_HPO_4_. The MgSO_4_ solution calls for 104 g/L of MgSO_4_.

### 2.2 Growing Mycelial Materials

The Δ*mpkA* mutant and the control were first grown on separate modified MAG-V plates for 5 days. Conidia from resulting conidial lawns were harvested using 5 mL of sterile DI water and a glass pipette. The harvested conidial suspension was placed into a conical tube, counted and diluted to the concentration of 8×10^6^ conidia/mL using modified YGV. The conical tube containing both the modified YGV and the conidia was vortexed, and 7.5 mL of the homogenized YGV-conidia mixture was added to a single 60×15 mm petri dish (Fisherbrand) to generate one mycelial material “disk.” This process was repeated to produce additional mycelial material disks. The filled petri dishes were placed into an incubator for 120 hours at 28□. The mycelial material formed on the top layer of liquid in each plate. Mycelial materials were harvested by transferring the entire contents of the petri dish to a beaker filled with diluted bleach, and then washed with DI water.

### 2.3 Growth Curve

To develop growth curves, 20 mycelial material petri dishes were prepared (section 2.2) for both the control and the Δ*mpkA* mutant. At each time point, the entire contents of two petri dishes for each strain were recovered via vacuum filtration (60 mm filter paper disk) and dried at 60□ until they reached constant weight. The average dry biomass of the two values at each time point was used to generate the growth curve.

### 2.4 Tensile Testing

Tensile testing was carried out on nominal 4 × 40 mm coupons cut (Universal Laser Systems VLS 3.6/6) from mycelial disks. At minimum, five coupons were cut from each disk (three disks per genotype). Before testing, coupons were hydrated in a 40:60 glycerol:water mixture for 1 minute [31], coated in petroleum jelly [32], and dried with warm air for approximately 1 minute on each side. Coupon dimensions were then precisely measured using a dissection microscope (Leica Microscope s9i Model meb115), and adhered to a paper frame (for stability) using cyanoacrylate adhesive (Loctite) and NaHCO_3_ powder (to accelerate drying [33]). The frame was mounted to the tensile testing device (Instron 3369) with double sided tape. Immediately prior to testing, the paper frame was cut on each side so only mycelial material was tensile tested. Two pieces of reflective tape were placed onto the paper frame to enable laser extensometer (Electronic Instrument Research Model LE-01) strain measurement during applied stress. Stress was measured by a sensor (Transducer Techniques Sensor) connected to the tensile testing device. All tests were stopped manually after material failure.

### 2.5 Mycelial Structure and Hyphal Morphology

For scanning electron microscopy (SEM) sample prep, fractured tensile-tested mycelial materials were removed from the paper frame, adhered to an aluminum stub using double sided carbon tape, sputter coated (Cressington 108 Manual Sputter Coater - Au/Pd target), and imaged using an FEI Nova NanoSEM 450. For transmission electron microscopy (TEM) sample prep, two mycelial disks from each genotype were cut into 1 × 1 cm squares and placed into a scintillation vial overnight at 4□. Vials were filled with a solution of 2% paraformaldehyde, 2.5% glutaraldehyde, in 0.1 M sodium cacodylate buffer at 7.4 pH. Samples were washed and submerged with fresh 0.1 M sodium cacodylate buffer, while being gently mixed for 10 minutes; this step was repeated twice. Samples were gently mixed in 0.1 M sodium cacodylate buffer and 50 mM glycine for 15 minutes. Samples were again washed and mixed with 0.1 M sodium cacodylate buffer for 10 minutes, three different times. Washed samples were incubated in 0.1 M sodium cacodylate buffer with 1% OsO4 for 1-2 hours, and rinsed with deionized water four times for 10 minutes each. The mycelial material was put through a dehydration series where each step was repeated three times for 10 minutes. Samples were first submerged in 35% ETOH, then 50% ETOH, 70% ETOH with 1% uranyl acetate, 70% ETOH, 95% ETOH, 100% ETOH, and 100% Acetone. Afterwards, samples were placed in 2:1 Acetone/Epon for 1 hour, then sectioned into 80 n slices using an ultramicrotome (Leica Microsystems UC7) and imaged on a FEI Tecnai T12 TEM. The diameter of the hyphae and thickness of the cell wall were measured at three different points (on the hyphae) from the TEM images of 20 different hyphae per genotype using ImageJ software.

### 2.6 Hyphal Resistance to Fragmentation

The mycelial disks (one disk per trial, four trials per genotype) were placed into a high-shear mixer (Hamilton Beach Model #58615) with 370 mL of water. The first sample (20 ml) was removed after 5 seconds, and subsequent samples (20 ml) were removed at 10s intervals. After collection, each sample was subjected to a particle size analysis (Malvern Mastersizer 3000) to provide a relative measure of average particle size, reported as average diameter (μm), as we have described previously [34]. Specifically, taking the 90th percentile of the particle size distribution (PSD) has been found to be representative of hyphal size.

### 2.7 Cell Wall Composition

The composition of rigid and mobile components was determined by analyzing peak volumes in 2D ^13^C-^13^ C spectra, employing 53 ms CORD and DP refocused J-INADEQUATE schemes, respectively. Solid-state NMR spectra were collected on an 800 MHz Bruker Avance Neo NMR spectrometer using a 3.2 mm HCN magic-angle spinning probe. ^13^C chemical shifts were calibrated on the tetramethylsilane (TMS) scale. Peak volumes were extracted using the integration function of Bruker Topspin software. To minimize uncertainties from spectral crowding, only well-resolved signals were considered for the compositional analysis. The identified NMR peaks, along with their resonance assignments and corresponding peak volumes, were visualized in a donut graph using Origin 2021.

Samples taken from four different disks per genotype were used to measure the percentage of melanin in a material. Mycelial materials were soaked in hexamethyldisilazane (HMDS) for 5 minutes, and air dried for at least 90 minutes (fume hood). The dried material was inserted into a cryogenic vial containing 3 glass beads (diameter 3 mm), and pulverized with a Mini Beadbeater (Biospec Products) at 5000 rpm for 1 minute. A small amount of the pulverized mycelial material was massed on a balance (g), recorded, and used in a commercial assay (Amplite® Fluorimetric Melanin Assay Kit, AAT Bioquest) to quantify melanin mass (g). The percentage of melanin from the measured mycelial material was calculated by dividing the mass of the melanin derived from the commercial kit by the mass of the dried mycelial material previously measured and used in the kit.

### 2.8 Asexual development (conidiation)

Fungal lawns were grown, using the same concentration and volume of spores to inoculate modified MAG-V plates. Conidia from these lawns were harvested using the same approach mentioned in section 2.2. A small aliquot of the conidial suspension was diluted by 1000x in water and 10 μL of the diluted conidial suspension was placed into a hemocytometer (Fisher Scientific), so the number of conidia could be counted under an inverted microscope (Zeiss Axiovert 200).

### 2.9 Germination and Total Branching Rate

Germination and total branching rates (the combined rate of hyphal germination and hyphal branching phenomena) were measured using a previously published protocol involving coverslips [25, 35]. Coverslips were sterilized while petri dishes filled with modified YGV media (without KCl) were placed into an incubator at 28□. Coverslips were submerged in Concanavalin A [35] for 20 minutes, rinsed with sterile DI water and dried under a flame for 1 hour. After 1 hour, conidia were harvested from plates using the same process mentioned in section 2.2, deposited into the modified YGV media at a concentration of 1×10^5^ conidia/mL, and added to each coverslip. Fresh 1 mL of conidial suspension was used for each coverslip. Coverslips were dried again under a flame for 1 hour before being added to the incubating petri dish. A coverslip was removed at every hour, starting from hour 9 until hour 16, and fixed in 3% formaldehyde (Macron Fine Chemicals) for 30 minutes minimum. 30 images of clearly defined and isolated hyphae at each time point were captured at 40x with a fluorescent microscope (Zeiss Axiovert 200) using calcofluor white (CFW; Sigma-Aldrich) stained coverslips. The number of germination tubes and number of branches from each of the 30 images were measured and recorded using ImageJ.

### 2.10 Hyphal Adhesion

Samples used for the hyphal adhesion assay were prepared by using a 2 mL growth medium with 3.6×10^6^ conidia/mL, harvested from a modified MAG-V plate, to fill four out of six wells in a 6well-plate (Thermo Scientific). The remaining two wells were filled with 2 mL uninoculated growth medium. Five μL of 2.0 μm diameter orange fluorescent beads (Sigma-Aldrich) were added to all wells before being incubated at 28□ for 20-21 hours. After incubation, the supernatent of the plate was emptied; the plate was submerged underwater, inverted, and gently agitated. 5 mL of calcofluor white was added to each well for 1 minute, then inserted into a plate reader first at 385 excitation wavelength - 470 emission wavelength. The fluorescence of each well was recorded (RFU), retested at 520 excitation wavelength -540 emission wavelength, and the fluorescence of each well was recorded again. For the control, the fluorescent intensity of hyphal beads was divided by the fluorescent intensity of calcofluor white. The hyphal adhesion was calculated by multiplying the previous flouorescent ratio by the average RFU of calcofluor white per gram of biomass in the control (calculated from preliminary studies). The degree of hyphal adhesion in the mutant was normalized using a calculation detailed in the supplementary information to allow for comparison to the control.

### 2.11 FTIR Spectroscopy

Samples for FTIR spectrometry were prepared by submerging a piece of mycelial material into 40:60 volume based glycerol:water mixture for 60 seconds and blotting the surface dry. One sample was inserted into the FTIR at a time, and scanned four times in the 4000 cm^-1^ to 650 cm^-1^ range using transmission mode in the FTIR software (PerkinElmer Spectrum). The peaks from the FTIR readings were analyzed via an infrared spectroscopy tables found in the literature [36].

### 2.12 Significance Testing

All quantitative data was subjected to T-tests to identify any significant differences between groups using Microsoft Excel.

## 3. RESULTS AND DISCUSSION

### 3.1 Mycelial Material Growth

For all assessments, mycelial materials were generated identically for the control (A1405) and the *mpkA* deletion mutant (Δ*mpkA*; A1404). The mycelial materials were grown as disks (**Figure 1A, B**), on the surface of static liquid growth medium at 28 °C for 120h. Compared to the control, the Δ*mpkA* mutant produced thinner, wetter-looking, material with less surface texture. **Figure 1C** shows material growth curves. For the first 75 hours, the control material grew exponentially at a rate of 0.41 h^-1^, while the Δ*mpkA* material grew at a rate of 0.43 h^-1^, with no significant difference between the two (*P* = 0.84). The material generated from the control stops increasing in mass after approximately 100 hours, but the Δ*mpkA* mutant material continued to increase in mass for 120 hours, leading to 50% more biomass (*P* = 0.001).

**Figure 1.**
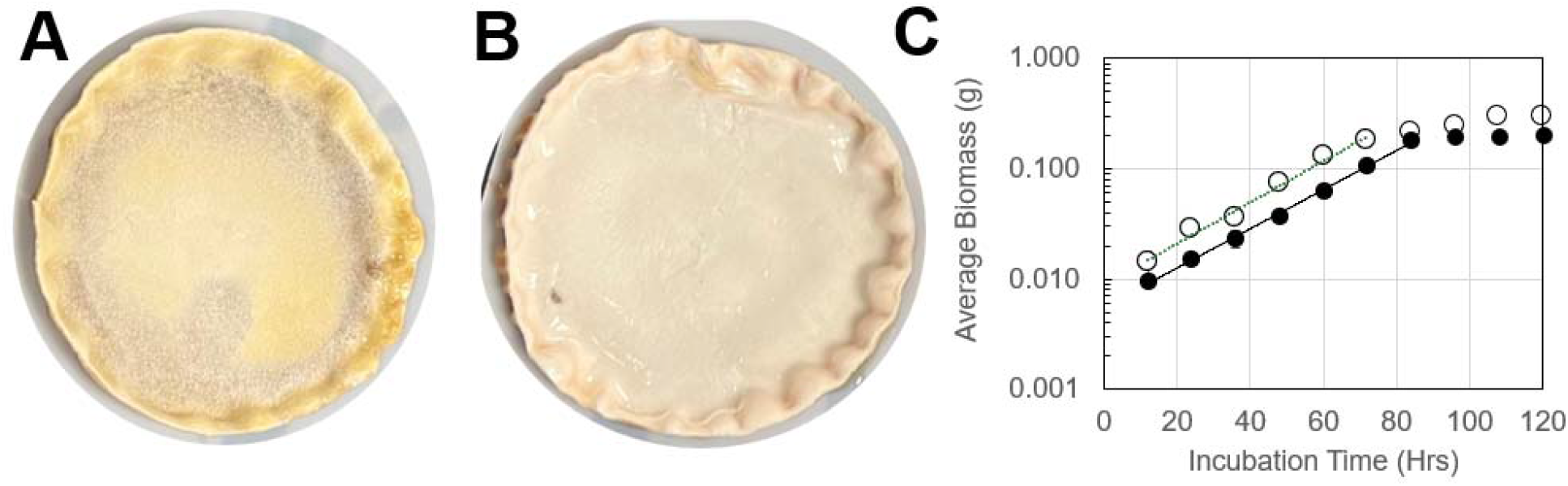
Mycelial material generation. Mycelial discs were grown from the (**A**) control (A1405) 5) and (**B**) Δ*mpkA* (A1404) fungal strains. (**C**) Average material biomass (g) for the control (•) and Δ*mpkA* (○) increased exponentially, with average specific growth rates (straight lines) of 0.41 and 0.43 h^-1^, and final biomass values of 0.20 and 0.30 g respectively. No significant difference in growth rates (*P* = 0.84); significant difference in final weights (*P* = 0.001). Error bars smaller than symbols.

### 3.2 Tensile Testing, Mechanical Properties, and Fracture Analysis

After 120 hours of growth, mycelial materials were carefully removed from the surface of the liquid growth medium, (prepared as described in the Methods section) and subjected to tensile testing. **Figure 2A** shows the strips used for testing, the paper frame on which the strips were mounted (**Figure 2B**), and one of these frames loaded onto the tensile testing apparatus (**Figure 2C**). Typical tensile-test curves for the control material and the Δ*mpkA* material are shown in **Figure 3**. For both genotypes, materials showed only linear elasticity before failure, a characteristic of brittle materials [37]. Data collected from the tensile test were used to determine: Young’s modulus (YM), ultimate tensile strength (UTS) and strain at failure (SF). Average values for these properties are shown in **Figure 4**. While both the control and the ΔmpkA material had similar YM (*P* = 0.449), material generated from the Δ*mpkA* mutant had significantly higher values of UTS (*P* = 0.003) and SF (*P* = 0.001) at 339 ± 18 kPa and 0.30 ± 0.02 respectively compared to the control which had a UTS of 213 ± 33 kPa and a SF at 0.22 ± 0.01. These results imply that the Δ*mpkA* material is stronger and more elastic than the control material.

**Figure 2.**
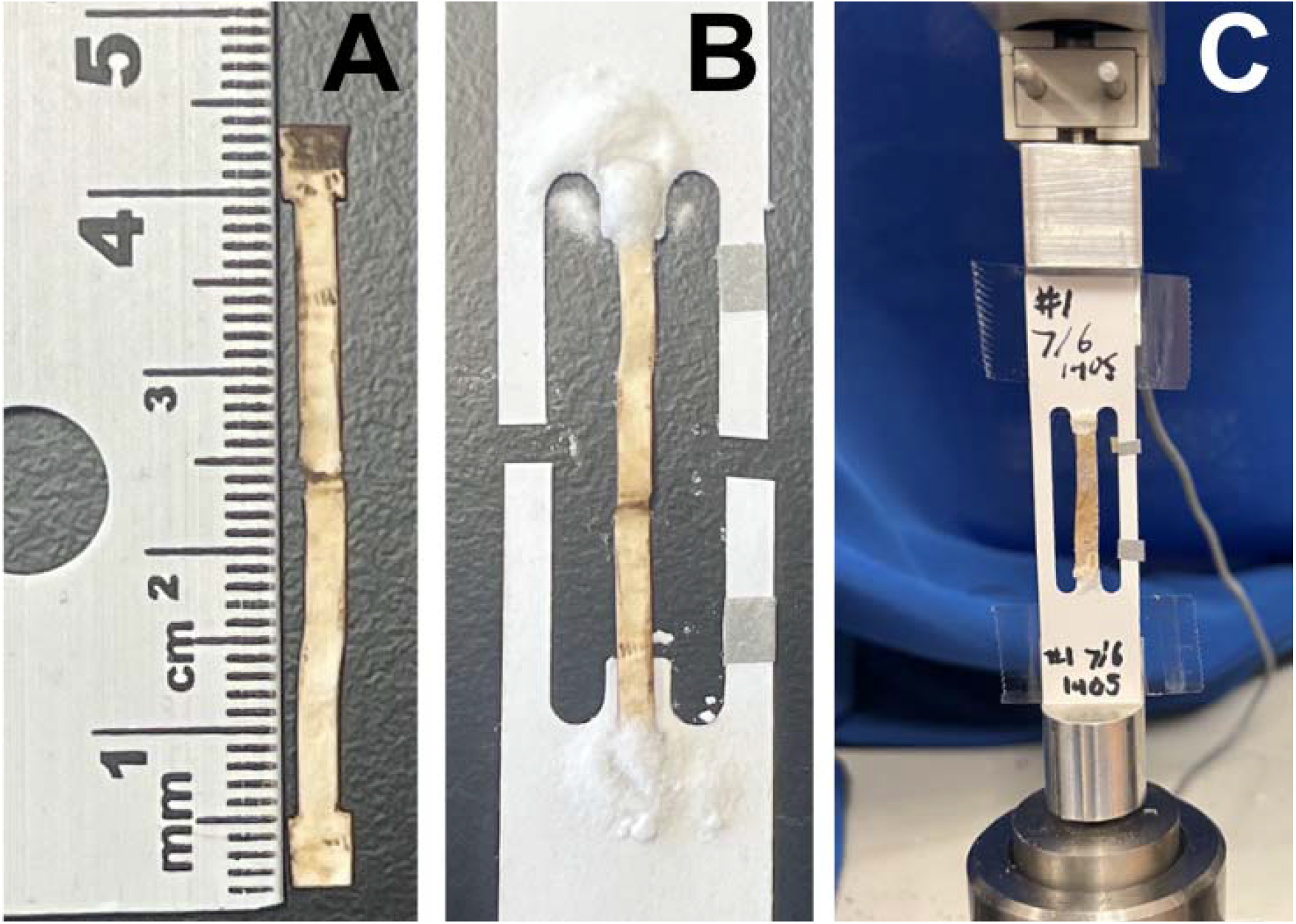
Tensile testing of mycelial material. Material discs were dried and then (**A**) cut into strips, which were (**B**) mounted in a paper frame which was then (**C**) mounted on load-frame for testing. Details in Methods.

**Figure 3.**
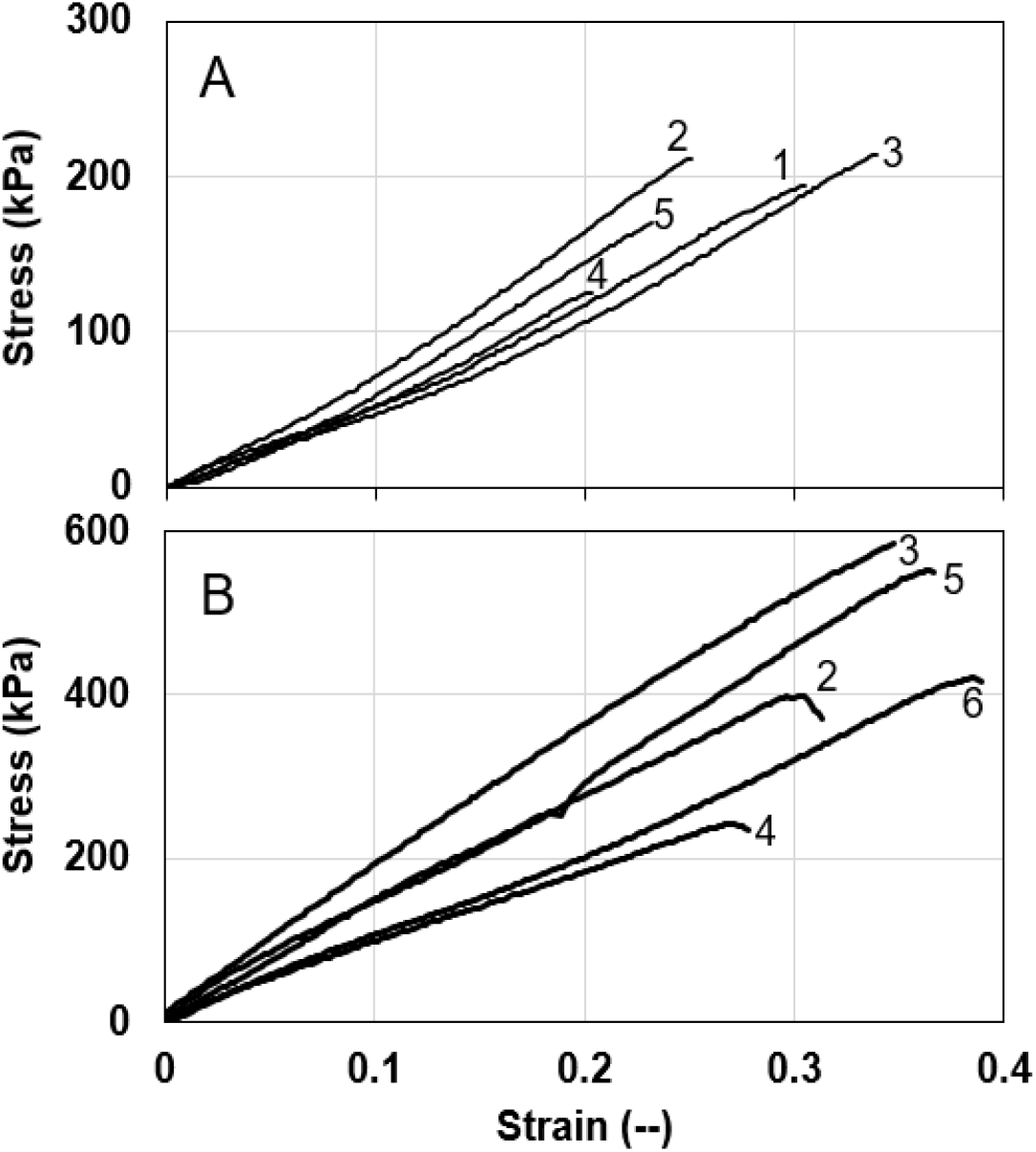
Typical stress-strain curves for mycelial material generated from the (**A**) control (A1405) and (**B**) Δ*mpkA* (A1404) fungal mutant. In both cases, five test strips were cut from the center of a single disc of mycelial material. Numbers on each graph indicate relative position of strips. For both fungal genotypes, the material shows linear elasticity before failure.

**Figure 4.**
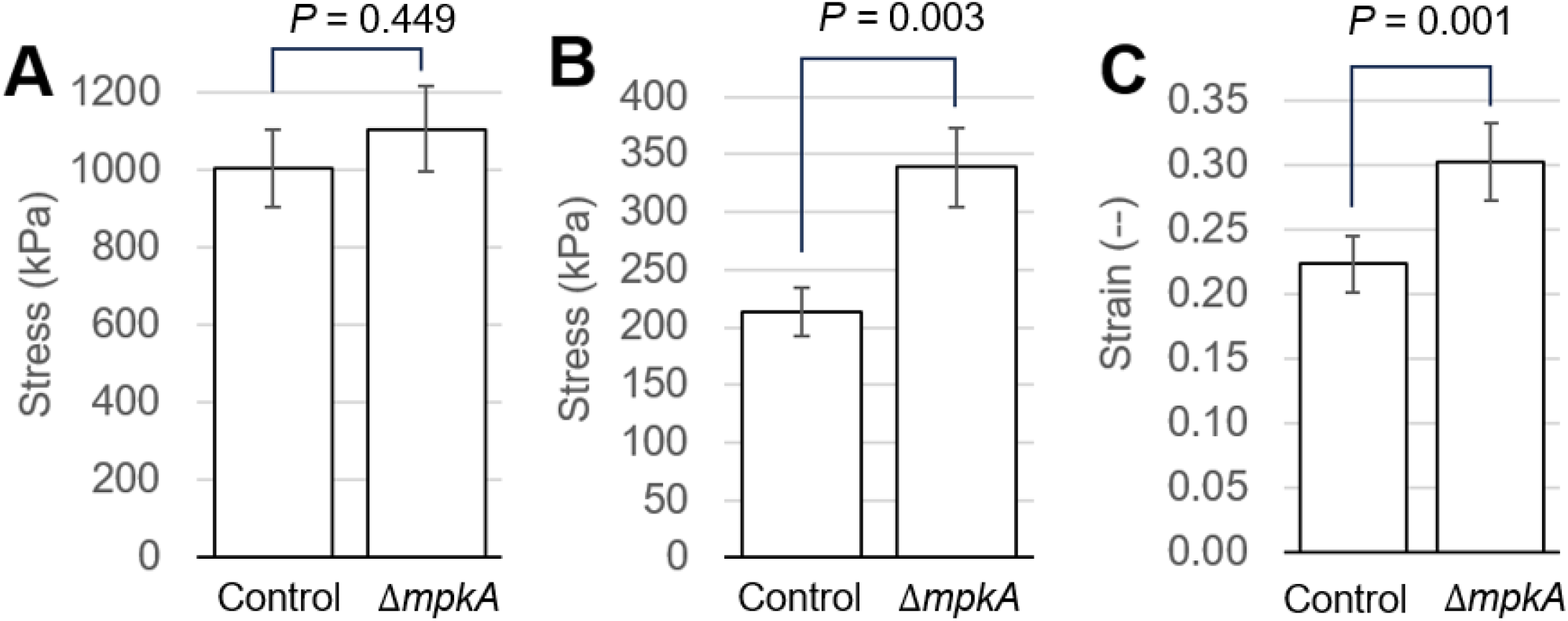
Mechanical properties determined from stress-strain testing (*n* = 15) of material generated from control (A1405) and *ΔmpkA* (A1404) mutant. (**A**) Average Young’s modulus, (**B**) ultimate tensile strength and (**C**) strain at failure.

### 3.3 Surface Morphology, Fracture Analysis, and Internal Morphology

To help understand the results from tensile testing, we used scanning electron microscopy (SEM) to analyze material surface morphology and morphology at the point of fracture. This allowed observation of the material’s three-dimensional structure, which was assessed for the degree of density (e.g. presence and/or size of void, hyphal packing) and morphological features. In general, compared to the control, we found the Δ*mpkA* material appeared to have (i) fewer developmental structures (i.e. conidia and Hülle cells.) and (ii) a more densely packed morphology with fewer visible voids (**Figure 5A,C)**. Similarly, the fracture point of the Δ*mpkA* material showed relatively more hyphae and fewer developmental structures. (**Figure 5B, D**). A previous study has shown that higher material density is correlated with increased mechanical strength [21], and the same appears to be true here. We suspect the lack of voids limits the number of unsupported positions that could potentially weaken the material.

**Figure 5.**
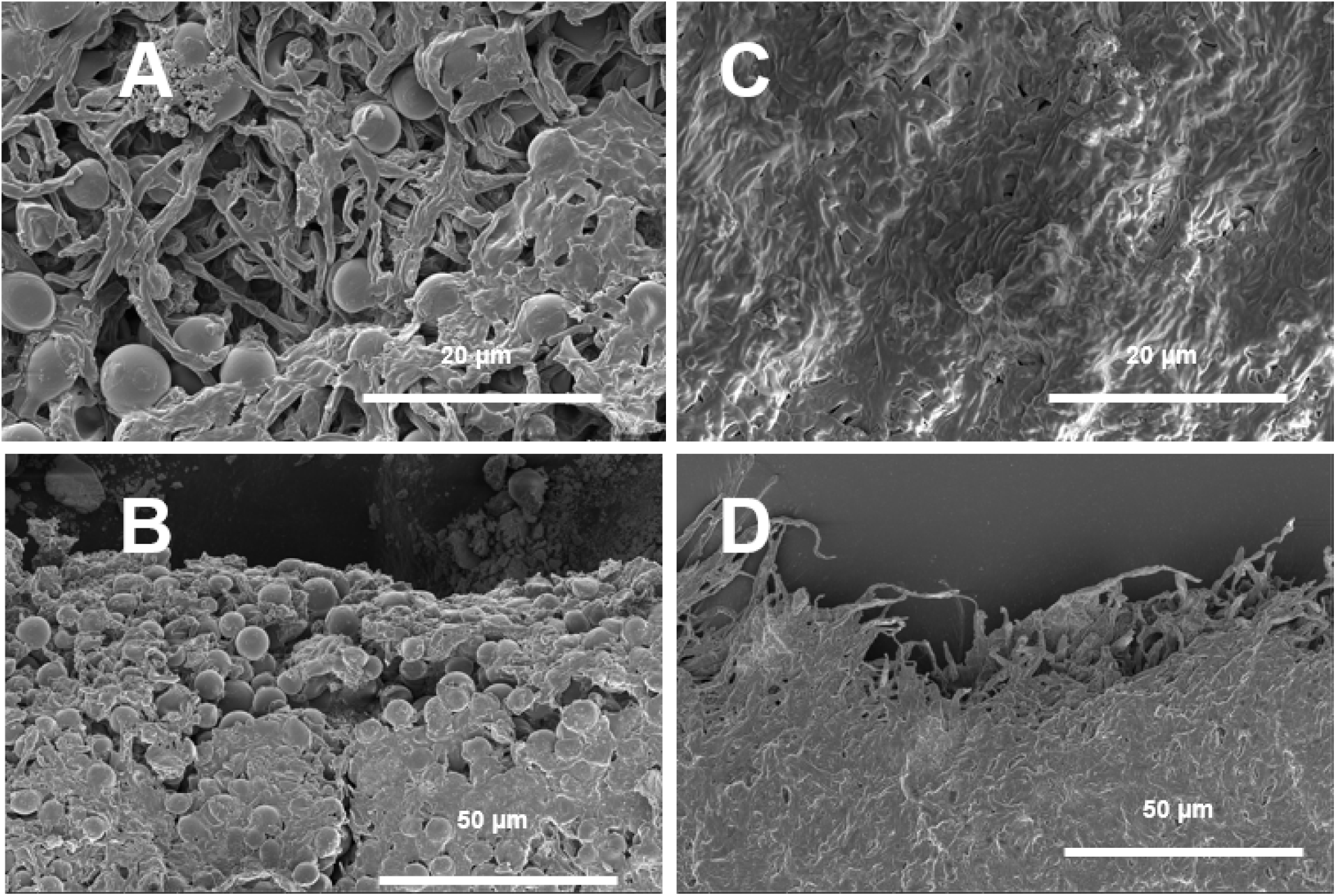
Representative SEM images of mycelial material three-dimensional structure and morphological features. (**A**) Control (A1405) sample surface showing an abundance of developmental structures (e.g., conidia, Hulle cells), loose packing of hyphae, and voids in the material. (**B**) Fracture point of the same control sample. (**C**) The Δ*mpkA* (A1404) sample surface ce shows fewer developmental structures, denser hyphal packing, and an absence of voids. (**D**) Fracture point of the same Δ*mpkA* sample.

To assess internal hyphal morphology, samples from both the control and the Δ*mpkA* mutant were analyzed via transmission electron microscopy (TEM, **Figure 6)**. From these images, hyphal cross-sections were used to determine average hyphal diameter (**Figure 6C**) and the average cell wall thickness (**Figure 6D**). We hypothesize that mycelial materials with larger hyphal diameters and thicker cell walls would be more equipped to handle larger forces, resulting in stronger materials. Compared to the control, Δ*mpkA* mutant hyphae showed a 50% larger hyphal diameter at 2124 ± 86 nm compared to a hyphal diameter of 1405 ± 70 nm in the control (*P* < 0.01); there was no significant difference in cell wall thickness.

**Figure 6.**
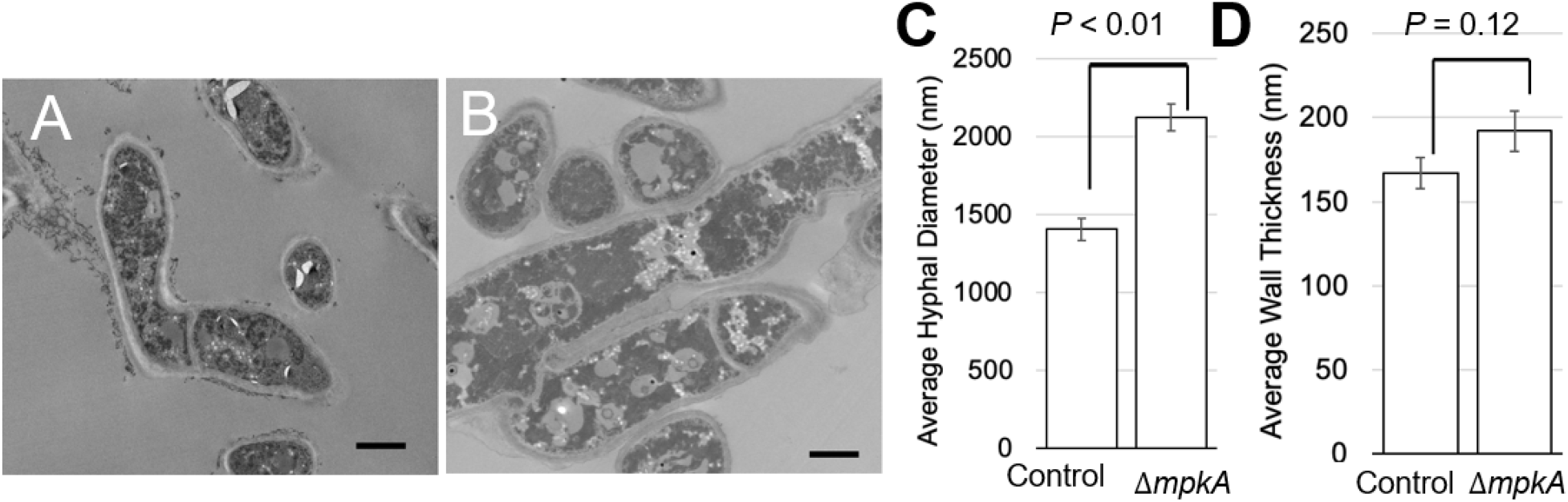
Representative TEM images showing the hyphal morphology of (**A**) control (A1405) and (**B**) Δ*mpkA* (A1404) mutant. Size bars = 1 μm. Measurements of hyphal morphology show that the (**C**) average hyphal diameter (nm) is significantly wider in the mutant and (**D**) cell wall thickness (nm) is similar.

### 3.4 Shear Stress Tolerance and Specific Fragmentation Rate

To assess relative material strength we used a fragmentation assay that we have described previously for fungi grown in suspension culture [34]. In this case, we added an entire disc of mycelial material to DI water, and subjected the material to high-shear mixing (**Figure 7)**. The fragmentation process in the high-shear mixer has two phases: (i) the initial cutting phase and (ii) the cavitation fragmentation phase [38]. The first phase occurs quickly (≤ 5 seconds) as the blades of the high-shear mixer accelerate from rest. During the initial mixing of the material, we assume that some mechanical shearing occurred without the presence of cavitation forces - implying the material’s tolerance to mechanical shear stress [39]. **Figure 7B** shows the Δ*mpkA* mutant has a larger size after this first phase at 2231 ± 46 nm compared to 1794 ± 12 nm in the control. This finding is consistent with the finding of higher ultimate tensile strength for the Δ*mpkA* material. During the second phase, the mycelia themselves are breaking, and the slope of the curve can be used to determine specific fragmentation rate (μm/s) [34], where there is no significant difference between the two materials (**Figure 7C**).

**Figure 7.**
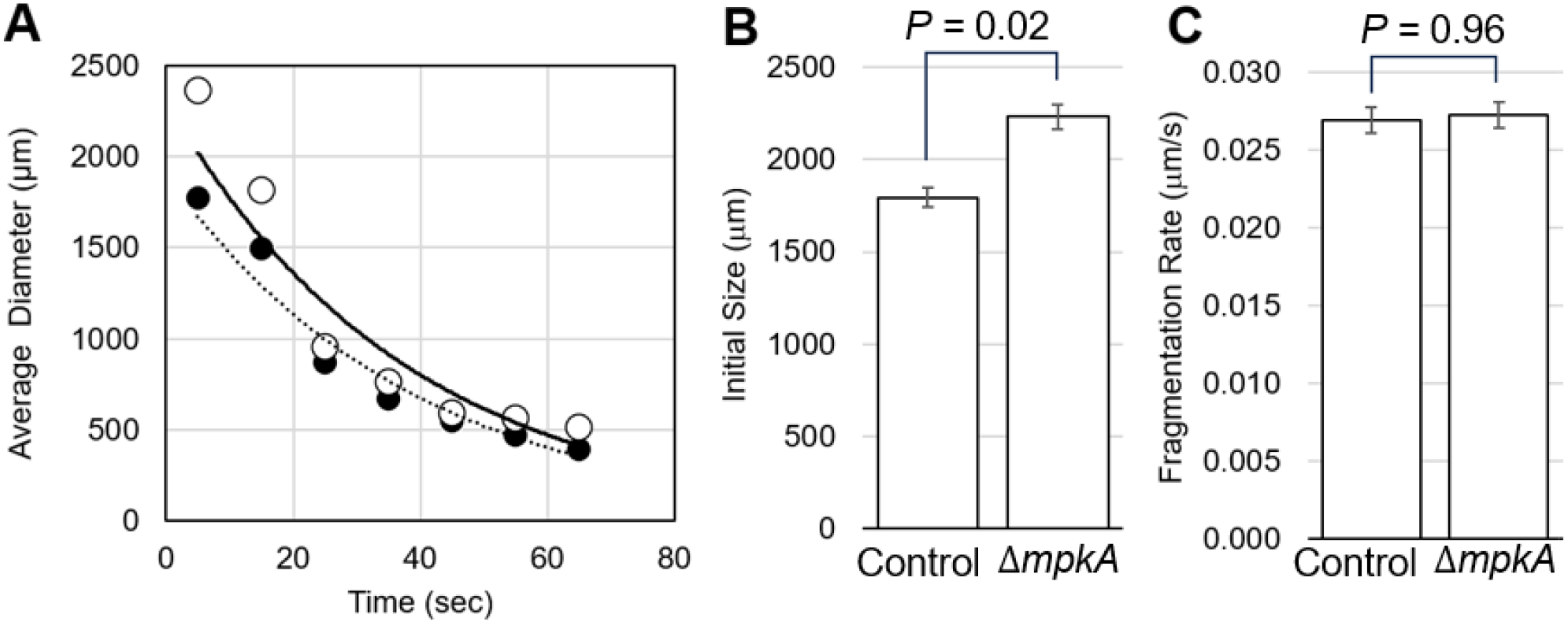
Relative material strength measured in a high-shear mixer. (**A**) Average size (n = 4) of fungal elements versus time in the high-shear mixer for material from the control (•) and Δ*mpkAA* (○) mutant. Lines indicate exponential fit used to determine specific fragmentation rate. (**B**) Average size of fungal elements for the control and *ΔmpkA* mycelial material after 5s of high-shear mixing are significantly different (*P* = 0.02). (**C**) Fragmentation rate, determined from exponential fits in A, are not significantly different (*P* = 0.96).

### 3.5 Cell Wall Composition

The filamentous fungal cell wall is essential for survival, providing protection from environmental, chemical, and mechanical stress. This structure is primarily composed of polysaccharides and melanin [22]. Within the mycelial cell wall of *Aspergillus* species, chitin physically associates with α-1,3-glucan and β-1,3-glucan to form rigid core structures, which are embedded within a soft, hydrated matrix composed of β-glucans with various linkages [40], including β-1,3, β-1,3/1,6, and β-1,3/1,4-linkages. This composite structure is further enveloped by an outer layer enriched with proteins, galactomannan, and galactosaminogalactan. Some of the polysaccharides are covalently bonded, such as chitin-β-1,3-glucan-galactomannan and chitin-β-1,3-glucan, thereby conferring resistance to alkali extraction [41].

To assess the cell wall composition of materials generated with both the control and Δ*mpkA* mutant, we used a high-resolution, solid-state NMR approach as has been described previously [42]. The results are shown in **Figures 8 and 9**. The 2D ^13^C-^13^C CORD experiment, which probes the rigid core of the fungal cell wall, revealed that it is composed of α-1,3-glucan, β-1,3-glucan, chitin and chitosan (**Figure 8 A,B**), consistent with the results reported in recent studies [43, 44]. The compositional analysis shows a significant decrease in α-1,3-glucan components in the Δ*mpkA* mutant (**Figure 8 A-C**), alongside a dramatic increase in the β-1,3glucan. Notably, no major changes were observed in the proportion of chitin and chitosan. In addition, employing 2D ^13^C-^13^C DP J INADEQUATE experiment, probed the mobile domain of the cell wall. We observed the alternation of galactomannan structure with selectively increased amount of α-1,2-mannose. Interestingly, the Δ*mpkA* mutant exhibited GalNAc, monomer of galactosaminogalactan (GAG) linkages at 101 ppm and 55 ppm which was absent in positive control. This may suggest the compensatory mechanism for the loss of α-1,3-glucan, which is crucial for aggregation and biofilm formation. Previous studies have shown that both α-glucan and GAG play a significant role in hyphal aggregation [45]. Our observations also indicate a significant loss of galactose in the mutant in comparison to positive control strain, which coincides with the emergence of GalNAc.

**Figure 8.**
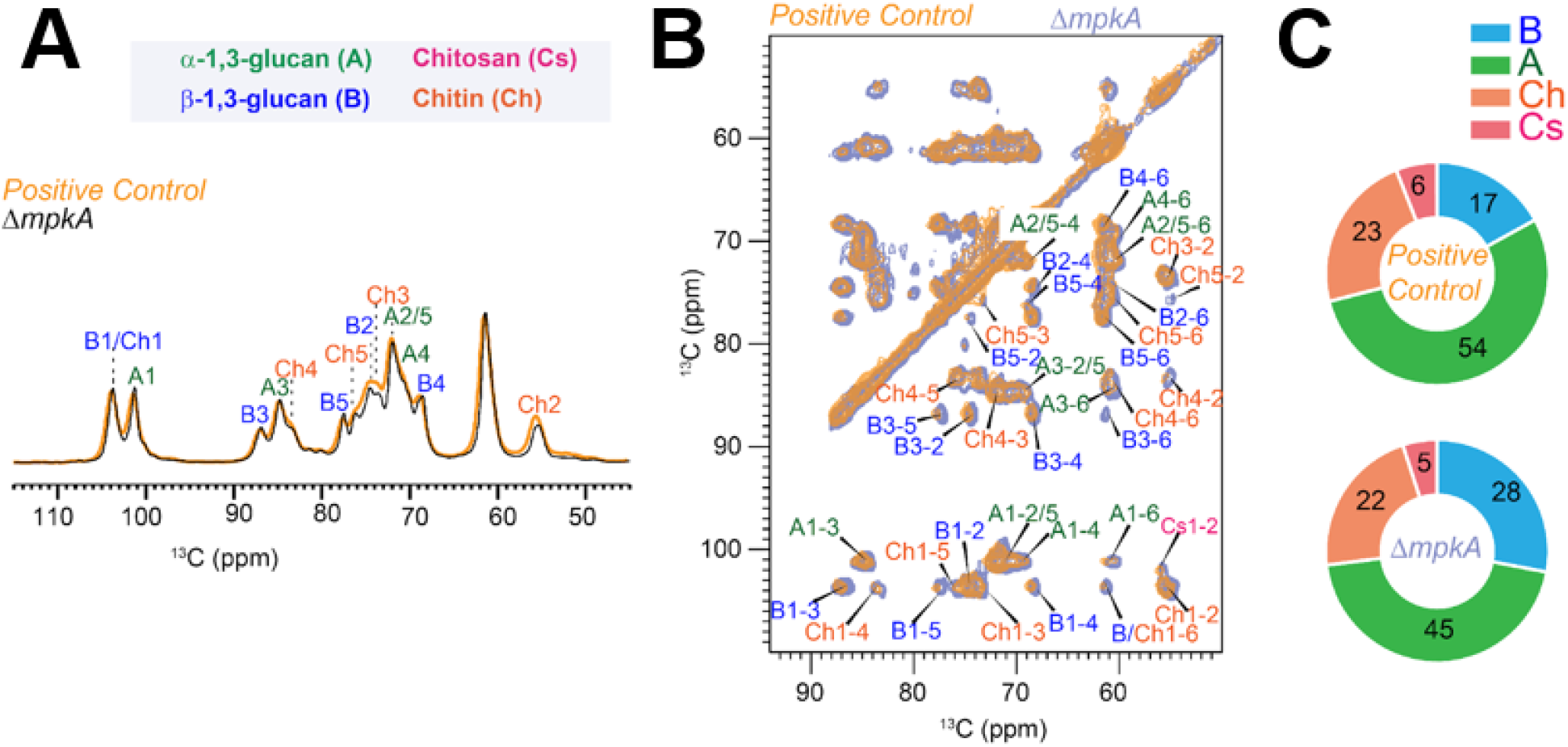
Composition of rigid polysaccharide probed by high-resolution ssNMR. (**A)** Overlay CP spectra of positive control and Δ*mpkA* mutant, with the rigid polysaccharide shown in orange and black spectra. The glucan types and its carbon numbers are abbreviated, and color coded as: α-1,3glucan (A, green), (β-1,3-glucan (B, blue), chitin (Ch, orange), chitosan (Cs, pink). (**B)** The 2D ^13^C-^13^C CORD, correlation explores the spatial arrangement of cross-linkages among α-1,3glucan, β-1,3-glucan, chitin, and chitosan. **(C)** The molar composition of these rigid components is determined by analyzing the peak volumes in the 2D ^13^C-^13^C CORD spectra.

**Figure 9.**
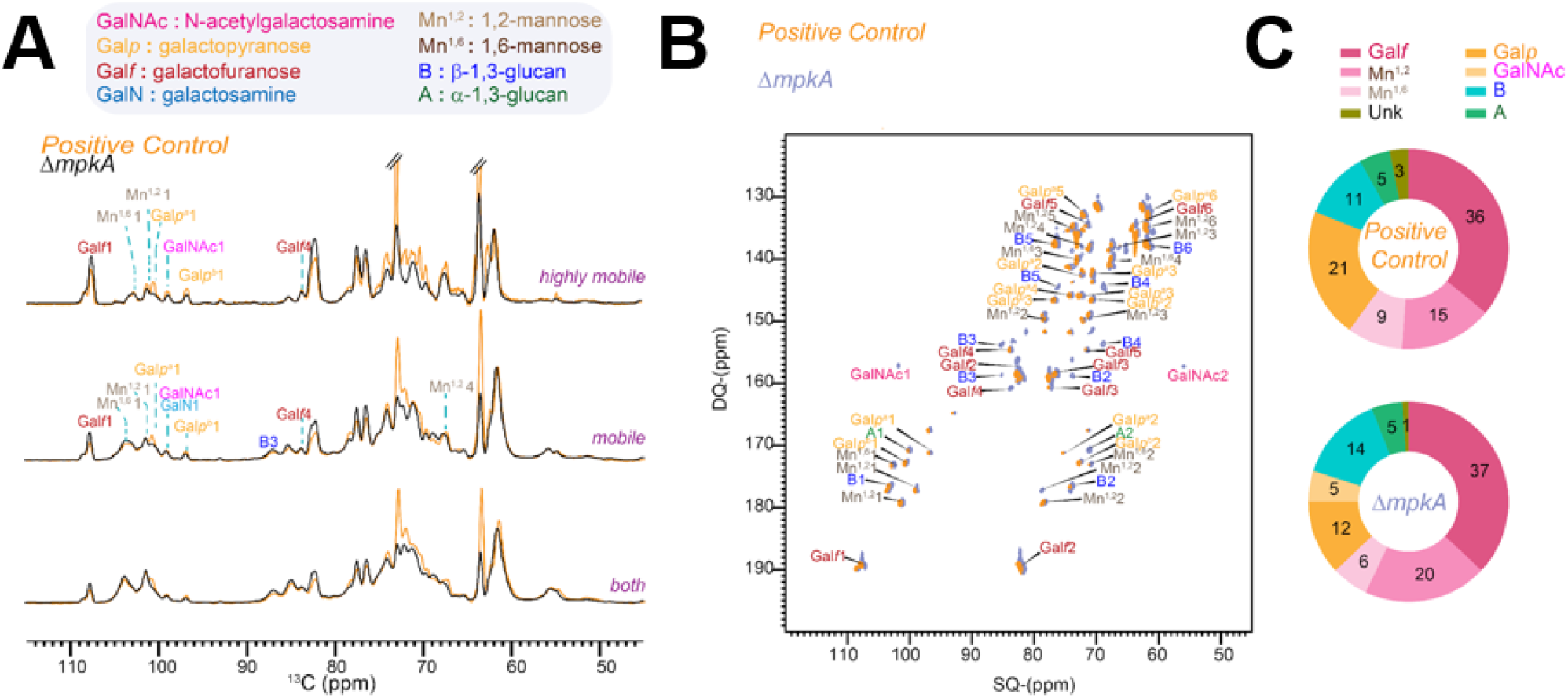
Composition of mobile carbohydrates. (**A)** Overlay of 1D spectra for the control (A1405) and Δ*mpkA* mutant (A1404), shown in orange and black, representing the highly mobile e region, semi-dynamic region, and cumulative region using INEPT, 2s DP, and 35s DP experiments. The glucan types, their monomers, and carbon numbers are labeled and color-coded ed as follows: galactofuranose (Gal*f*, pink), α-1,2-mannose (Mn^1,2^, light brown), α-1,6-mannose (Mn^1,6^, brown), galactopyranose (Gal*p*, yellow), and N-acetyl galactosamine (GalNAc, magenta). **(B)** The through-bond correlations 2D 1^13^C-^13^C -DP J INADEQUATE correlation spectrum, shown in orange and grey for positive control and mutant. **(C)** The molar composition of these mobile components determined by analyzing the peak volumes in the 2D DPINADEQUATE spectra. The GM molecules are color-coded in pink shades, while the GAG components are shaded in orange in the donut graph. The legend corresponds to the abbreviations, linking the monomers to their assignments in the INADEQUATE spectra.

Melanin plays a role in preserving and hardening the fungal cell wall. Here, the percentage of melanin found in the cell wall was determined using a commercial assay (**Figure 10**). Compared to the control, material generated from the Δ*mpkA* mutant contained nearly twice the amount of melanin (*P* = 0.02) per dry cell weight (DCW) at 0.16 ± 0.01% compared to the control which had 0.09 ± 0.01% melanin per DCW (**Figure 10**).

**Figure 10.**
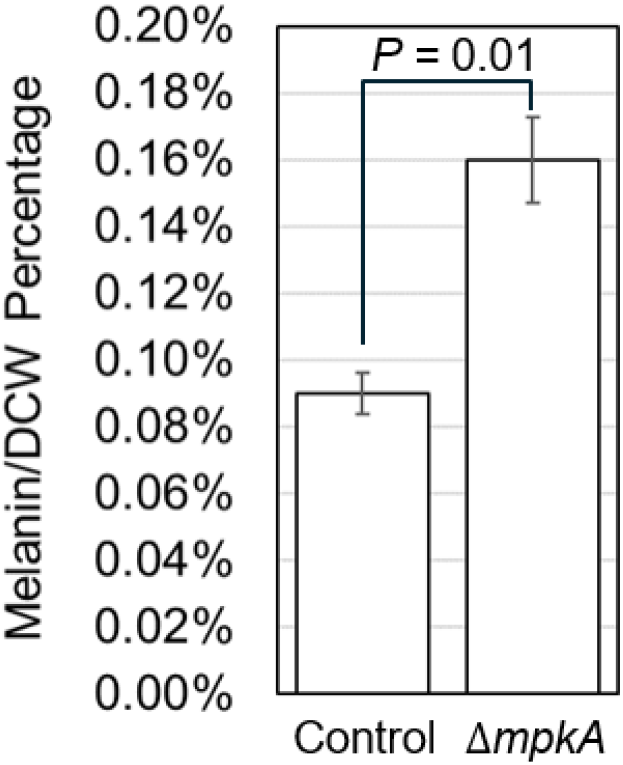
Cell wall melanin content. Compared to the control, the ΔmpkA mutant produces significantly more (∽80%) melanin (*P* < 0.01).

### 3.6 Asexual Development

The mechanical properties of mycelial materials are dependent on the interaction of hyphae within a grown material [13, 46, 47]. Hyphae extend and develop during vegetative growth, which is interrupted by asexual development [48-50]. SEM observations of the two genotypes used here implies that material generated with the Δ*mpkA* mutant contains fewer developmental structures. To better understand the extent of this phenomena, lawns of both the control and the mutant were grown on the surface of agar plates. Conidia from each genotype were collected and counted, and results are shown in **Figure 11**. Compared to the control, the Δ*mpkA* mutant produced 40% fewer conidia at 339 ± 12 conidia/mL compared to the control’s 544 ± 34 conidia/mL (*P* = 0.02).

**Figure 11.**
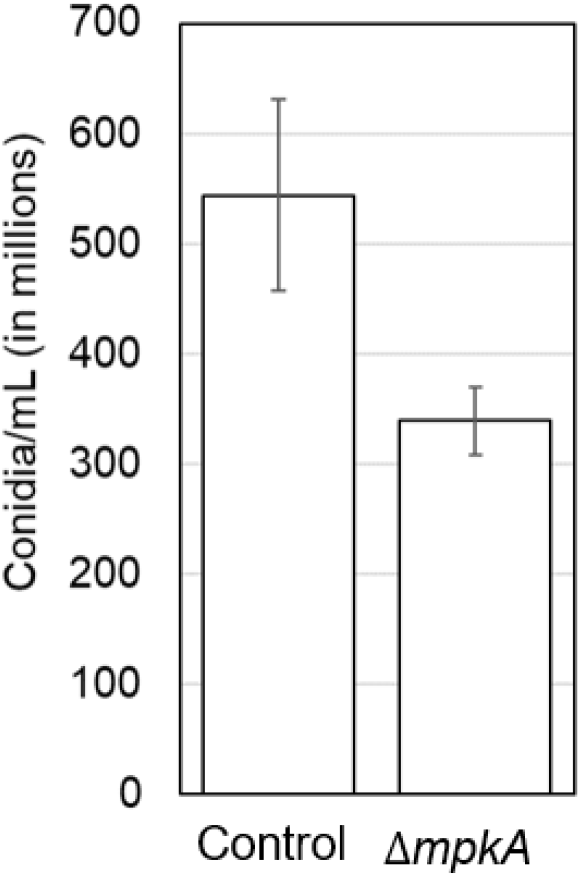
Conidia collected, show that compared the control the Δ*mpkA* produces significantly fewer (40%) conidia (*P* = 0.02).

### 3.7 Germination, and Total Branching Rates

To determine if there was a difference between the germination and branching rates of the two stains, we grew cells on cover slips as described previously [35]. This allows visualization of individual spores that develop into hyphae, thus allowing determination of germination and total branching rates. The control and Δ*mpkA* mutant displayed similar germination rates at 0.10 ± 0.02 germinated hyphae/hr and similar total branching rates at 0.11 ± 0.02 total branches/hr. The p value for each phenotype, germination rate (*P =* 0.46) and total branching rate (*P =* 0.52) indicates that there is no significant difference in either germination or total branching rates between the control and Δ*mpkA* mutant.

### 3.8 Hyphal Adhesion

Higher hyphal adhesion leads to a more developed biofilm that has the potential to contribute to altered mechanical properties [51]. The hyphal adhesion of the control and Δ*mpkA* material were 1.39×10^5^ ± 0.24 x10^5^ and 1.10 x10^5^ ± 0.22 x10^5^ respectively. The p value between the genotypes (*P =* 0.40) reveals there is no significant difference in hyphal adhesiveness.

### 3.9 FTIR Results

Testing via fourier transform infrared (FTIR) spectroscopy provides the ability to characterize the water content of heterogenous natural materials, and others have used this approach to characterize mycelial materials [17, 36]. In particular, FTIR is able to identify water found in a material through OH bonds, which are detected in the wavelength range from 3600 cm^-1^ to 3200 cm^-1^ [52, 53]. **Figure 12** shows typical FTIR traces for the Δ*mpkA* mutant and the control, with a clear difference for the peaks within the wavelengths 3600 cm^-1^ to 3200 cm^-1^. This height difference indicates a noticeable increase in the volume of water found in the Δ*mpkA* mutant compared to the amount of water found in the control. This finding is consistent with results from a previous study [21], in which the mutant with higher ultimate tensile strength and strain at failure, also contained more water, implying a correlation between increased water capacity and increased ultimate tensile strength and strain at failure.

**Figure 12.**
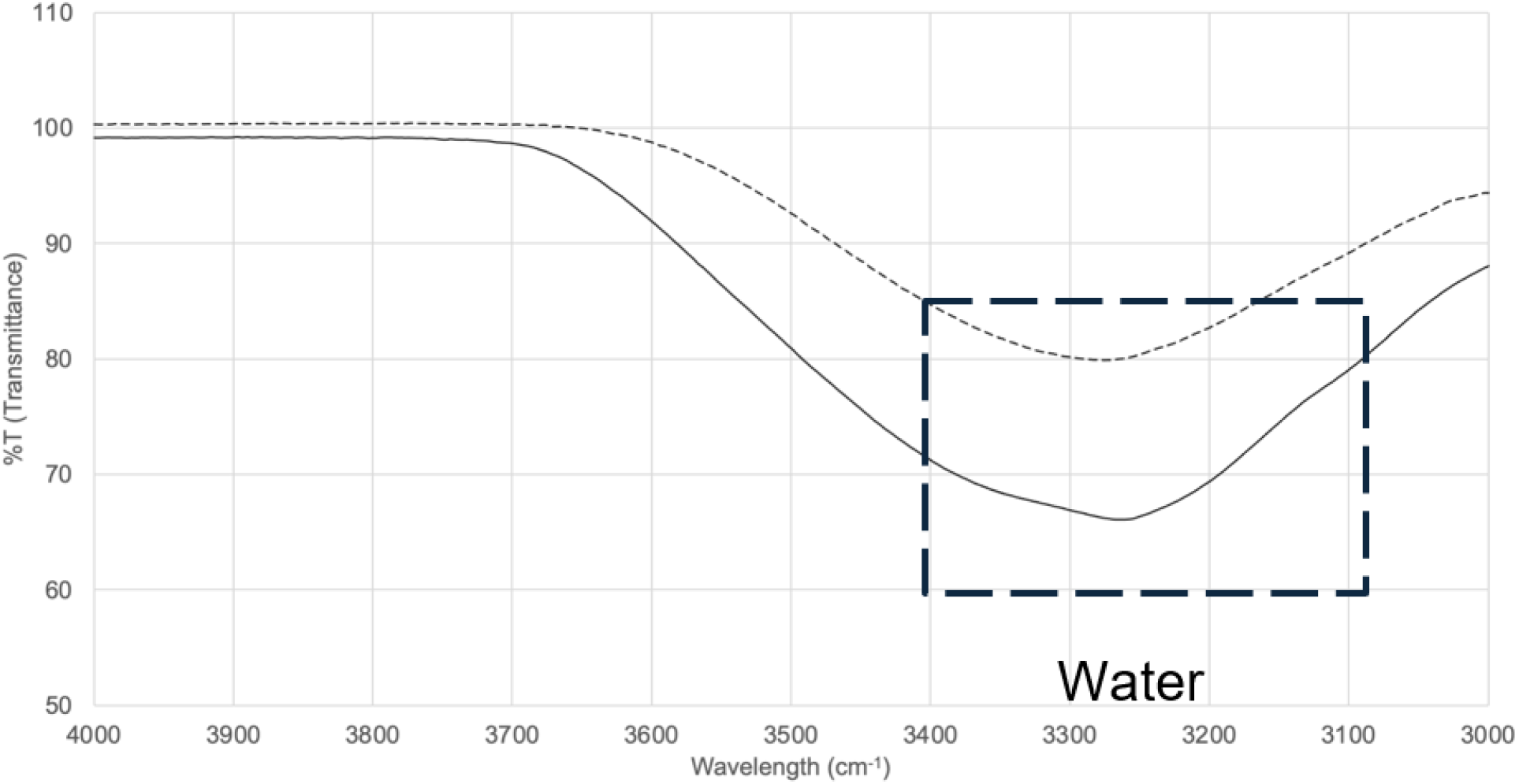
FTIR traces for material grown from control (- - -) and Δ*mpkA* (––) mycelial materials. Annotated graph showing the difference in water between the control and the Δ*mpkA mutant with a close up image highlighting additional traces*. This graph also suggests that there might be additional chemical differences found between the control and the Δ*mpkA* mutant.

## 4. Conclusion

Here we compared two mycelial materials – one made exclusively from an *A. nidulans* control strain and the other from an isogenic, deletion-mutant lacking the last kinase in the CWI signaling pathway, MpkA. We found the deletion of the *mpkA* gene leads to materials with higher ultimate tensile strength, strain at failure, and resistance to shear-stress. This result was obtained through both the tensile testing of the actual material, and a novel fragmentation assay [34].

To determine possible causes for this behavior, we characterized a number of different phenotypes of both the individual fungal strains and the resultant mycelial material. We found that, compared to the control, material generated from the Δ*mpkA* mutant had a significantly higher total amount of biomass. Consistent with this, SEM images show a reduction in the number of developmental structures, and growth on plates confirmed the Δ*mpkA* mutant produces significantly fewer developmental structures. This reduction appears to have allowed the mycelia to be more closely packed (i.e., fewer apparent voids in SEM images) both in general and at the point of fracture during tensile testing. In addition, FTIR results imply the Δ*mpkA* strain has a larger amount of water. These behaviors are consistent with a previous study which tested mycelial material generated from a different species of filamentous fungi (*Schizophyllum commune*) with a deletion of a surface hydrophobin gene (Δ*sc3*) [21]. Similar to our work, they found that (compared to an isogenic control) material generated by the deletion strain had increased density (more closely packed mycelia) and a higher amount of water, which led to increased ultimate tensile strength, and higher strain at failure [21]. In work here, the Δ*mpkA* mutant also has significantly larger hyphae, which contain more melanin and more β-glucan than the control. These differences may have also played a role in the significant differences in material properties. Together, these findings imply that it is possible to use genetic means to control fungal phenotype, leading to significant differences in resultant mycelial-material mechanical properties. Additional studies will be required to develop a deep understanding of how material mechanical properties can be rationally manipulated using various genetic approaches.

## Acknowledgements

We gratefully acknowledge significant assistance from personnel in the UMBC, Keith R. Porter Imaging Facility regarding SEM preparation and imaging; Drs. Michael Duffy and Joao Santos (Engineering Testing, LLC) for their assistance and guidance with material tensile testing; Dr. Govind Rao (UMBC, Center for Advanced Sensor Technology) for use of the laser cutter to prepare samples; Dr. Jorge Almodovar for general guidance with this project. We also thank the Electron Microscopy Core Imaging Facility at the University of Maryland, School of Medicine for their critical role with the entire TEM imaging process. Finally, we thank the UMBC, Molecular Characterization and Analysis Complex for use of the FTIR instrument.

## REFERENCES

1. Ramesh, M., K. Palanikumar, and K.H. Reddy, Plant fibre based bio-composites: Sustainable and renewable green materials. Renewable and Sustainable Energy Reviews, 2017. 79: p. 558–584.

2. Nakhate, P. and Y. Van Der Meer, A systematic review on seaweed functionality: a sustainable bio-based material. Sustainability, 2021. 13(11): p. 6174.

3. Olivetti, E.A. and J.M. Cullen, Toward a sustainable materials system. Science, 2018. 360(6396): p. 1396–1398.

4. Gawel, E., N. Pannicke, and N. Hagemann, A path transition towards a bioeconomy— The crucial role of sustainability. Sustainability, 2019. 11(11): p. 3005.

5. Nations, F.a.A.O.o.t.U., Bioeconomy for a sustainable future. 2021, Food and Agriculture Organization of the United Nations.

6. Babu, K.M., Silk from silkworms and spiders as high-performance fibers, in Structure and Properties of High-Performance Fibers. 2017, Elsevier. p. 327–366.

7. Verma, N., S.E. Jujjavarapu, and C. Mahapatra, Green sustainable biocomposites: Substitute to plastics with innovative fungal mycelium based biomaterial. Journal of Environmental Chemical Engineering, 2023. 11(5): p. 110396.

8. Abhijith, R., A. Ashok, and C.R. Rejeesh, Sustainable packaging applications from mycelium to substitute polystyrene: a review. Materials today: proceedings, 2018. 5(1): p. 2139–2145.

9. Meyer, V., et al., Growing a circular economy with fungal biotechnology: a white paper. Fungal biology and biotechnology, 2020. 7: p. 5–5.

10. Angelova, G.V., M.S. Brazkova, and A.I. Krastanov, Renewable mycelium based composite – sustainable approach for lignocellulose waste recovery and alternative to synthetic materials – a review. 2021. 76(11-12): p. 431–442.

11. Girometta, C., et al., Physico-mechanical and thermodynamic properties of mycelium-based biocomposites: a review. Sustainability, 2019. 11(1): p. 281.

12. Alaneme, K.K., et al., Mycelium based composites: A review of their bio-fabrication procedures, material properties and potential for green building and construction applications. Alexandria Engineering Journal, 2023. 83: p. 234–250.

13. Vandelook, S., et al., Current state and future prospects of pure mycelium materials. Fungal biology and biotechnology, 2021. 8(1): p. 1–10.

14. Whabi, V., B. Yu, and J. Xu, From Nature to Design: Tailoring Pure Mycelial Materials for the Needs of Tomorrow. Journal of Fungi, 2024. 10(3): p. 183.

15. Li, K., et al., Engineered living materials grown from programmable Aspergillus niger mycelial pellets. Materials Today Bio, 2023. 19: p. 100545.

16. Jo, C., et al., Unlocking the magic in mycelium: Using synthetic biology to optimize filamentous fungi for biomanufacturing and sustainability. Materials Today Bio, 2023. 19: p. 100560.

17. Haneef, M., et al., Advanced materials from fungal mycelium: fabrication and tuning of physical properties. Scientific reports, 2017. 7(1): p. 41292.

18. Deeg, K., et al., Greener Solutions: Improving performance of mycelium-based leather. Final Report to MycoWorks, 2017: p. 1–24.

19. Appels, F.V., et al., Fungal mycelium classified in different material families based on glycerol treatment. Communications biology, 2020. 3(1): p. 1–5.

20. Schaak, D., Bio-Manufacturing Process (Patent# 16/363052), in United States Patent and Trademark Office, U.S.P.a.T. Office, Editor. 2019, Ecovative Design LLC.: United States of America.

21. Appels, F.V.W., et al., Hydrophobin gene deletion and environmental growth conditions impact mechanical properties of mycelium by affecting the density of the material. Scientific reports, 2018. 8(1): p. 4703.

22. Gow, N.A.R., J.-P. Latge, and C.A. Munro, The fungal cell wall: structure, biosynthesis, and function. Microbiology spectrum, 2017. 5(3): p. 10–1128.

23. Levin, D.E., Cell wall integrity signaling in Saccharomyces cerevisiae. Microbiology and molecular biology reviews, 2005. 69(2): p. 262–291.

24. Fujioka, T., et al., MpkA-dependent and-independent cell wall integrity signaling in Aspergillus nidulans. Eukaryotic cell, 2007. 6(8): p. 1497–1510.

25. Chelius, C.L., et al., Phosphoproteomic and transcriptomic analyses reveal multiple functions for Aspergillus nidulans MpkA independent of cell wall stress. Fungal Genetics and Biology, 2019. 125: p. 1–12.

26. De Souza, C.P., et al., Functional analysis of the Aspergillus nidulans kinome. PLoS One, 2013. 8(3): p. e58008.

27. McCluskey, K., The fungal genetics stock center. from molds to molecules, 2003. 52: p. 245–262.

28. Nayak, T., et al., A versatile and efficient gene-targeting system for Aspergillus nidulans. Genetics, 2006. 172(3): p. 1557–1566.

29. Harris, S.D., J.L. Morrell, and J.E. Hamer, Identification and characterization of Aspergillus nidulans mutants defective in cytokinesis. Genetics, 1994. 136(2): p. 517–532.

30. Käfer, E., Meiotic and mitotic recombination in Aspergillus and its chromosomal aberrations. Advances in genetics, 1977. 19: p. 33–131.

31. Sethi, A., et al., Moisturizers: the slippery road. Indian journal of dermatology, 2016. 61(3): p. 279–287.

32. Hamishehkar, H., et al., A comparative histological study on the skin occlusion performance of a cream made of solid lipid nanoparticles and Vaseline. Research in pharmaceutical sciences, 2015. 10(5): p. 378–387.

33. Katti, D. and N. Krishnamurti, Anionic polymerization of alkyl cyanoacrylates: In vitro model studies for in vivo applications. Journal of applied polymer science, 1999. 74(2): p. 336–344.

34. Quintanilla, D., et al., A fast and simple method to estimate relative, hyphal tensilestrength of filamentous fungi used to assess the effect of autophagy. Biotechnology and bioengineering, 2018. 115(3): p. 597–605.

35. Chelius, C., et al., Dynamic transcriptomic and phosphoproteomic analysis during cell wall stress in Aspergillus nidulans. Molecular & Cellular Proteomics, 2020. 19(8): p. 1310–1329.

36. Lecellier, A., et al., Differentiation and identification of filamentous fungi by high-throughput FTIR spectroscopic analysis of mycelia. International journal of food microbiology, 2014. 168: p. 32–41.

37. Wilkerson, R.P., Biomimetic” Nacre-Like”, Metal-Compliant-Phase Ceramics Produced via Coextrusion. 2018: University of California, Berkeley.

38. Vancleef, A., et al. Reducing the Induction Time Using Ultrasound and High-Shear Mixing in a Continuous Crystallization Process. Crystals, 2018. 8, DOI: 10.3390/cryst8080326.

39. Shirgaonkar, I.Z., R.R. Lothe, and A.B. Pandit, Comments on the mechanism of microbial cell disruption in highpressure and highspeed devices. Biotechnology Progress, 1998. 14(4): p. 657–660.

40. Latgé, J.-P. and T. Wang, Modern biophysics redefines our understanding of fungal cell wall structure, complexity, and dynamics. Mbio, 2022. 13(3): p. e01145–22.

41. Latgé, J.-P. and G. Chamilos, Aspergillus fumigatus and Aspergillosis in 2019. Clinical microbiology reviews, 2019. 33(1): p. 10–1128.

42. Chakraborty, A., et al., A molecular vision of fungal cell wall organization by functional genomics and solid-state NMR. Nature communications, 2021. 12(1): p. 6346.

43. Kang, X., et al., Molecular architecture of fungal cell walls revealed by solid-state NMR. Nature communications, 2018. 9(1): p. 2747.

44. Dickwella Widanage, M.C., et al., Adaptative survival of Aspergillus fumigatus to echinocandins arises from cell wall remodeling beyond β− 1, 3-glucan synthesis inhibition. Nature Communications, 2024. 15(1): p. 6382.

45. Miyazawa, K., et al., Both galactosaminogalactan and α-1, 3-glucan contribute to aggregation of Aspergillus oryzae hyphae in liquid culture. Frontiers in Microbiology, 2019. 10: p. 2090.

46. Nguyen, P.Q., et al., Engineered living materials: prospects and challenges for using biological systems to direct the assembly of smart materials. Advanced Materials, 2018. 30(19): p. 1704847.

47. Alemu, D., M. Tafesse, and A.K. Mondal, Mycelium-based composite: The future sustainable biomaterial. International journal of biomaterials, 2022. 2022(1): p. 8401528.

48. Adams, T.H. and W.E. Timberlake, Developmental repression of growth and gene expression in Aspergillus. Proceedings of the National Academy of Sciences, 1990. 87(14): p. 5405–5409.

49. Adams, T.H., J.K. Wieser, and J.-H. Yu, Asexual sporulation in Aspergillus nidulans. Microbiology and molecular biology reviews, 1998. 62(1): p. 35–54.

50. Mirabito, P.M., T.H. Adams, and W.E. Timberlake, Interactions of three sequentially expressed genes control temporal and spatial specificity in Aspergillus development. Cell, 1989. 57(5): p. 859–868.

51. Gow, N.A.R. and M.D. Lenardon, Architecture of the dynamic fungal cell wall. Nature Reviews Microbiology, 2023. 21(4): p. 248–259.

52. Perakis, F., et al., Vibrational spectroscopy and dynamics of water. Chemical reviews, 2016. 116(13): p. 7590–7607.

53. Eisenberg, D. and W. Kauzmann, The structure and properties of water. 2005: Oxford University Press, USA.

