## Supplementary Figure 1 for "*Aspergillus nidulans* cell wall integrity kinase, MpkA, impacts cellular phenotypes that alter mycelial-material mechanical properties"

Abs = Absorbance

${Normalized Fluorescence}_{\Delta mpkA}\left( \frac{g}{RFU} \right)=(\left〖 \left( Abs) \right〗_{\Delta mpkA,CFW}-{(Abs)}_{blank,CFW} \right)\times\left( \frac{{grams}_{chitin}}{{Abs}_{fluorescence}} \right)\times\left( \frac{{grams}_{hyphal biomass}}{{grams}_{chitin}} \right)\times\left( \frac{{hyphal area growth rate}_{control}}{{hyphal area growth rate}_{\Delta mpkA}} \right)\times\left( \frac{{CFW bonding affinty}_{control}}{{CFW bonding affinity}_{\Delta mpkA}} \right)\times\left( \frac{{biomass growth rate}_{control}}{{biomass growth rate}_{\Delta mpkA}} \right)$

$\frac{Adhesion}{g}= \frac{{Fluorescence}_{beads}}{Normalized Fluorescence}=\frac{RFU}{g/{RFU}}=\frac{Adhesion (a ratio of fluorescence)}{g}$

Previous studies where the fluorescence of massed hyphae stained with calcofluor white were conducted. In these studies samples were grown in shake flasks where exclusively vegetative growth occurs. These studies revealed the ratios of:

1) ${{Abs}_{fluorescence}:grams}_{chitin}$

and

2) ${{grams}_{chitin}:grams}_{hyphal biomass}$

Supplementary Figure 1. Equation for normalizing ∆*mpkA* mutant fluorescence in adhesion assay.
